# Development of a flexible 3D printed TPU-PVC microfluidic devices for organ-on-a-chip applications

**DOI:** 10.1101/2024.12.20.629528

**Authors:** Rodi Kado Abdalkader, Satoshi Konishi, Takuya Fujita

## Abstract

The development of cost-effective, flexible, and scalable microfluidic devices is crucial for advancing organ-on-a-chip (OoC) technology for drug discovery and disease modeling applications. In this study, we present a novel 3D-printed flexible microfluidic device (3D- FlexTPU-MFD) fabricated through a one-step fused deposition modeling (FDM) process using thermoplastic polyurethane (TPU) as the printing filament and polyvinyl chloride (PVC) as the bonding substrate. The device’s compatibility was evaluated with various cell types, including human primary myoblasts, human primary endothelial cells (HUVEC), and human iPSC-derived optic vesicle (OV) organoids. Myoblasts cultured within the device exhibited high viability, successful differentiation, and the formation of aligned myotube bundles, outperforming conventional well-plate cultures. Additionally, iPSC-derived OV organoids- maintained viability, displayed neurite outgrowth, and sustained expression of the eye marker PAX6. These results demonstrate that the 3D-FlexTPU-MFD effectively supports cell growth, differentiation, and alignment, making it a promising platform for tissue modeling and OoC applications in future.

## 1. Introduction

In the field of drug discovery and disease modeling, there is a growing recognition of the limitations associated with traditional animal models and classical *in vitro* systems^1^. While these methods have been invaluable, they often fall short in replicating the complex physiological environments found in humans. As a result, there is a pressing need for alternative models that can provide more accurate and relevant data, ultimately improving the drug development process.

Micro physiological systems (MPS) or Organ-on-chip (OoC) technology has emerged as one such promising alternative^2^. These models are designed to recapitulate the microarchitecture and functions of human organs by integrating living cells into a three-dimensional (3D), dynamic environment in microfluidic devices^3^.

To advance the development of these systems, it is crucial to employ fast, simple, and cost-effective fabrication technologies. However, current OoC fabrication methods often rely on micro-/ nano-fabrication techniques like photolithography and PDMS casting, which involve multiple steps and require expensive materials and instruments in a clean room condition^4,5^. Recently, 3D printing, particularly with stereolithography (SLA) or digital light processing (DLP) printers using photoresins, has been used to fabricate microfluidic devices in less time and costs^5,6^. However, the use of organic solvents during the cleaning process and the cytotoxic nature of some resins can make these devices less suitable for cell culturing of fragile cells such as primary human cells and human induced pluripotent stem cells (iPSC) derived cells^7,8^.

We previously succeeded in fabricating micropatterned microfluidic devices using fused deposition modeling (FDM) with polyethylene terephthalate glycol (PETG) clear filaments, where murine myoblast cells could be cultured in an aligned manner^9^. However, the fused polymer lines can easily scatter light, presenting challenges during imaging and cell observation. Additionally, common polymers like polylactic acid (PLA) and PETG struggle to bond effectively to clear substrates such as glass, or plastic clear polymers such as polyvinyl chloride (PVC) without the addition of gluing materials, which complicate the process and can introduce toxic elements to the cells.

To overcome these challenges, we opted for thermoplastic polyurethane (TPU) polymer as the filament and the clear PVC as the bonding substrate for the filament deposition. TPU offers several advantages, including flexibility, biocompatibility, and ease of fabrication, making it a cost-effective alternative to more sophisticated and time-consuming methods^10,11^. More importantly, TPU possesses physicochemical properties that can enable bonding though hydrophobic interactions between TPU and PVC, as well as inter-crosslinking between both polymers thereby enhancing the bonding process^12^.

In this study, we have developed a 3D printed flexible TPU microfluidic device (3D- FlexTPU-MFD) that enables the culturing of human primary myoblast cells and their differentiation as well as human iPSC-derived optical vesicles (OV) organoids and tested their functions and characteristics. This model represents a significant step forward in creating more design-flexible, resource affordable and reliable OoC prototype models for drug applications purposes.

## 2. Materials and Methods

### 2.1. Device fabrication

3D-FlexTPU-MFD was fabricated by using FDM 3D-printing techniques (Anycubic technology) provided with 1.75 mm translucent clear TPU filaments (Saintsmart). Designs were issued by tinker cad (https://www.tinkercad.com) (**Supplemental figure 1**). Then STL files of the designs were sliced and processed using Ultimaker Cura software considering the following parameters: layers height: 0.01-0.02 mm and shell thickness: 2-10 layers. TPU filaments were then directly deposited on clear A4 size PVC sheet of 1 mm thickness. After printing the edges were trimmed and stored at room temperature. The fidelity of the printing was assessed by measuring the width of printing channel with using imageJ software acorrding the equation:

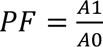

In this equation, PF, A1, and A0 represent printing fidelity, the channel’s actual widths, and the channel’s theoritical width. Scanning electron microscopy (SEM) analysis was performed using KEYENCE (VE-9800) for structure presentation of printed constructs. To test liquid flow in the microfluidic channels, a solution of sodium fluorescen (Nacalai) in water (1 mg mL^-1^) was employed.

### 2.2. Culturing of cells in 3D-FlexTPU-MFD

Microfluidic devices were sterilized by UV irradiation for 60 minutes, after which the channels were coated with matrigel (Thermo Fisher Scientific, Inc., Waltham, MA, USA) for 60 minutes in a humidified incubator at 37°C with 5% CO2.

Human primary normal skeletal myoblasts (HSkM) (Gibco) were cultured in DMEM (Wako, Osaka, Japan) medium, supplemented with 2% horse serum (Thermo Fisher Scientific), to induce cell differentiation. Cells were seeded at a density of 4.8 × 10⁴ per channel or per well (96-well plate) and incubated for 24 hours at 37°C with 5% CO2. The culture medium was replaced daily with fresh medium.

For culturing HUVEC cells, channels were pre-coated with matrigel as mentioned above. Then, 1×10⁴ cells were seeded in DMEM supplemented with 10 % v/v FBS, 10 ng mL^-1^ bFGF (Wako, Osaka, Japan) and 1ng mL^-1^ EGF (Wako, Osaka, Japan). After 24 hours, the medium was replaced, and the culture medium was refreshed daily.

Human iPSCs, 585A1^13^, was procured from Riken Cell Bank and handled in accordance with the guidelines stipulated by the ethics committees of Ritsumeikan University (Approved number 2021-004-3). Culture dishes were pre-coated with imatrix-511 (Nippi) overnight at 37°C. 1×10^4^ viable cells in mTeSR^TM^ plus medium supplemented with 10 µM Y27632 were seeded in 6-well plate cells. The next day cell culture medium was changed to mTeSR^TM^ plus only. The culture medium was refreshed with fresh mTeSR^TM^ plus medium every two days. Subsequently, embryoid bodies (EBs) were produced by adding 0.5-1×10^5^ viable cells in mTeSR^TM^ plus medium supplemented with 10 µM Y27632 in 10 cm ultra-low attachment dishes (Sumitomo, Tokyo, Japan). After three days, the mTeSR^TM^ plus medium was replaced with the chemical induction medium, consisting of fresh E6 medium (Thermo Fisher Scientific, Inc., Waltham, MA, USA) supplemented with 10 µM TGF-β kinase/activin receptor-like kinase inhibitor (SB-505124) (Santa Cruz), 10 µM Y27632 and BMP4 (20 ng mL^-1^). These conditions were maintained for 3 days. EBs were then transferred into 6-well plates coated with matrigel (6-8 EBs per well). Subsequently, the medium was switched to E6 medium supplemented with 10 µM Y27632 and 500 nM retinoic acid (Wako, Osaka, Japan) for 7 days. On day 20, OV organoids aggregates were picked with 27 G syringe needle and 1-2 clusters in E6 medium supplemented with 10 µM Y27632 were inserted per a channel of the device. The culture medium was refreshed with E6 medium supplemented with 10 µM Y27632 twice a day. For the determination of neurite outgrowth, brightfield images were employed and the length of neuron growth at different days were measured using imageJ software.

### 2.3. Determination of cell viability

Cell viability was assessed by the live staining with Calcein AM (Dojindo Molecular Technologies, Inc.). Briefly, cells were incubated with Calcein AM at a final concentration of 10 μg mL^-1^ in HBSS+ buffer at 37°C for 60 minutes. The cells were then washed once with HBSS+ and subjected to fluorescence microscopic imaging (KEYENCE, Tokyo, Japan).

### 2.4. Structure analysis and immunofluorescence staining

Cells were fixed with 4% paraformaldehyde in PBS for 20 minutes at 25°C. Subsequently, cells were permeabilized with 0.5% Triton X-100 in PBS for 10 minutes. Next, cells were incubated with ActinGreen 488 or ActinRed 555 (Thermo Fisher Scientific) and Hoechst 33342 (Dojindo Molecular Technologies, Inc.) in a final concentration of 10 µg mL^-1^ at 25 °C for 60 minutes. For performing immunofluorescence, following permeabilization, cells were blocked with a blocking buffer (5% bovine serum albumin, 0.1% Tween-20) at 25°C for 60 minutes. Next, cells were incubated with primary antibodies diluted in the blocking buffer (anti-PAX6; Proteintech Group, Inc.) for overnight at 4°C. After washing several times, the cells were incubated at 25°C for 60 minutes with the appropriate secondary antibody. Imaging was performed using fluorescence microscopy (KEYENCE, Tokyo, Japan) and confocal laser scanning microscope FV3000 (OLYMPUS) where z-scans of 2-3 µm were obtained. Images were then analysed using FV31S-SW software. For the investigation of cell morphology, ImageJ software was employed (National Institute of Health, Maryland, USA). For the determination of myotubes orientation, the Directionality function analysis as well as OrientationJ ImageJ.

### 2.5. Data visualization and statistics

Data are represented as mean ± S.E.M of at least three tests. The unpaired t-test and Tukey’s test were performed using GraphPad prism 8 (GraphPad Software, La Jolla California, USA). Data mining and visualization was conducted by Orange 3 software (Version 3.23.1; Bioinformatics Laboratory, Faculty of Computer and Information Science, University of Ljubljana, Slovenia, Python Jupyter notebook 6.1.4, with Pandas and Bioinfokit packages, and matplotlib and seaborn packages.

## 3. Results

### 3.1. Fabrication of the 3D-FlexTPU-MFD

The fabrication process utilized fused deposition modeling (FDM)-based 3D printing, in which TPU filaments were deposited onto a PVC substrate, resulting in strong polymer fusion and bonding (**Figure 1a**). Following the deposition of TPU onto PVC, a robust bond was formed, capable of withstanding a pulling force of up to 1 kg without failure (**Supplemental Figure 2**). To evaluate the structural integrity and printing fidelity, three microfluidic channel designs were tested: a letter-shaped channel (“R”), four straight channels (W1, W2, W3 and W4), and a serpentine channel (**Figure 1b, i–iii**). The printed devices demonstrated well-defined channel structures with consistent boundaries. High-resolution images of the serpentine channels filled with sodium fluorescein revealed smooth channel cross-sections, confirming successful fabrication (**Figure 1b, iv**) (**Supplemental video**). Fluorescence imaging of sodium fluorescein-filled channels W1 (15mm length, 0.5 mm width), W2 (15mm length, 1 mm width), and W3 (15mm length, 2 mm width) was performed to assess channel uniformity (**Figure 1c, i**). Fluorescence intensity profiles measured across the channel width showed uniform distribution of the dye, indicating consistent channel formation (**Figure 1c, ii**). Quantitative analysis of the printing fidelity ratio revealed notable differences among the channels (**Figure 1c, iii**). W2 and W3 achieved a fidelity ratio close to 1, indicating excellent dimensional accuracy and structural reproducibility. In contrast, W1 had a fidelity ratio slightly below 1, suggesting minor deviations from the intended design. This makes W2 and W3 more suitable for applications requiring high fidelity and consistent channel dimensions.

**Figure 1.**
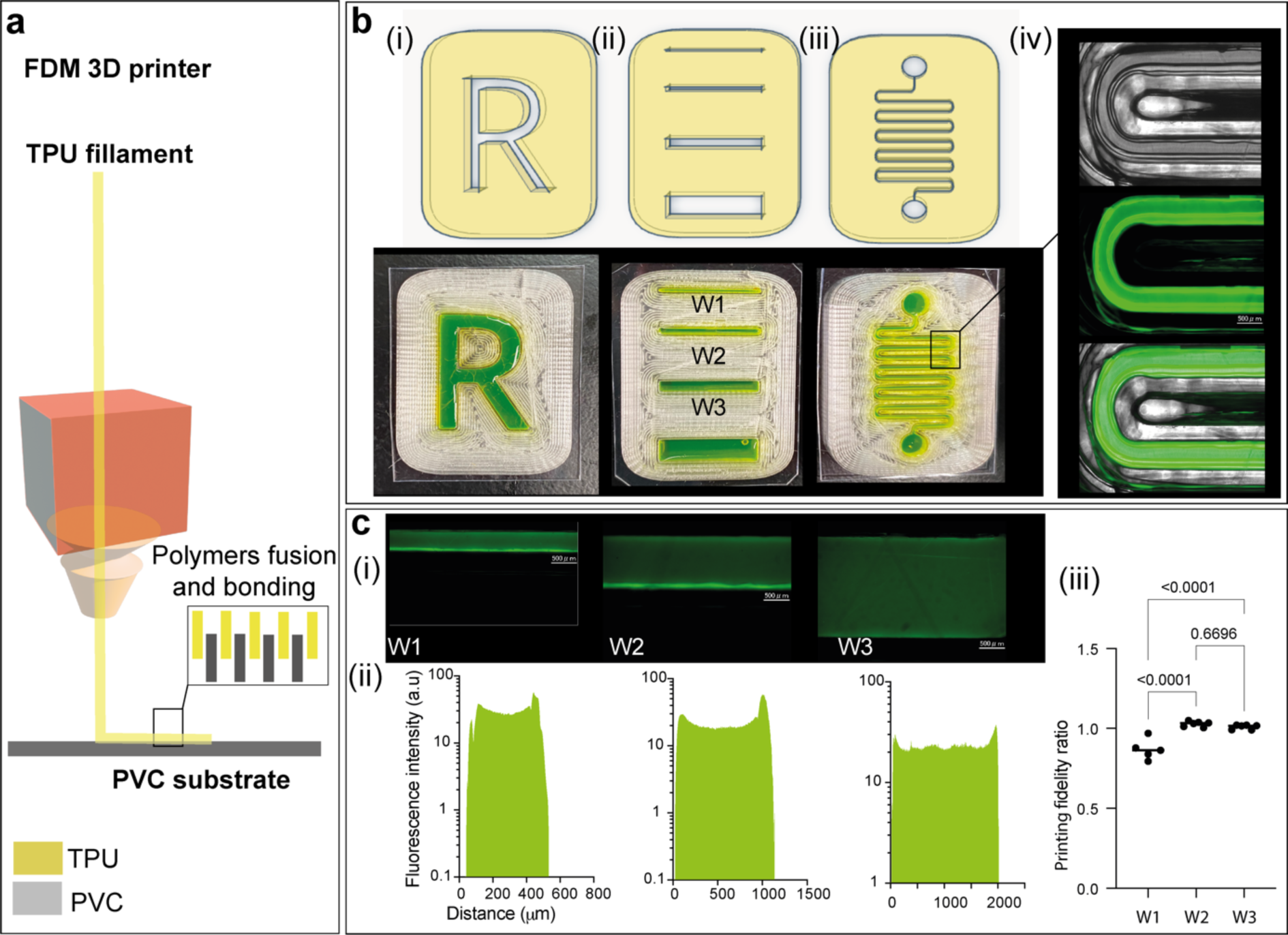
Design, fabrication, and characterization of the open top 3D-FlexTPU microfluidic devices. (a) Schematic illustration of the fabrication process for the 3D-FlexTPU microfluidic device using fused deposition modeling (FDM) 3D printing. TPU filaments are printed onto a PVC substrate, enabling fusion and bonding of the polymers. (b) Microfluidic channel design and fidelity assessment: (i–iii) Schematic diagrams of the designs (top) and corresponding results of printed microfluidic devices (bottom), including: (i) Letter “R” shape, (ii) Rectangular straight channels (W1, W2, W3), (iii) Serpentine channel geometry. (iv) High-resolution images showing the sectional structure and fidelity of serpentine microfluidic channels filled with sodium fluorescein. Scale bar: 500 μm. **(c)** Evaluation of channel fidelity: (i) Fluorescence images of sodium fluorescein-filled channels W1, W2, and W3. Scale bars: 500 μm. (ii) Fluorescence intensity profiles along the width of channels W1, W2, and W3, confirming consistent channel formation. (iii) Printing fidelity ratio for the three channel designs. (p < 0.0001) for W1 and W3 vs to W2.

Building on these findings, we designed a 3D-printed flexible microfluidic device (3D- FlexTPU-MFD) for cell culture applications using the same FDM-based fabrication method (**Figure 2a, i and ii**). The fabrication process involves several sequential steps. First, TPU polymers were deposited onto PVC sheets to form microfluidic channels, inlets, and outlet reservoirs (**Figure 2b, i**). Next, PVC covers are applied to seal the channels (**Figure 2b, ii**), followed by trimming of the device edges to finalize the fabrication (**Figure 2b, iii**). An actual image of the completed 3D-FlexTPU-MFD is presented in **Figure 2c, i**, demonstrating the actual printing of microfluidic channels and reservoirs, which include three chambers. The elasticity of the device shown through multiple cycles of manual twisting, as shown in **Figure 2c, ii**. Despite repeated flexing, the device maintained its structural integrity, confirming its durability and suitability for dynamic applications. Moreover, a cross-sectional scanning electron microscopy (SEM) image (**Figure 2c, iii**) shows the layered structure of TPU adhered to the PVC substrate, clearly depicting the cell chamber and inlet areas with robust bonding to the PVC. We then tested the 3D-FlexTPU-MFD with HUVEC cells and observed strong adherence, as indicated by the flattened morphology of the cells visualized through actin staining on PVC surfaces. Additionally, the cells exhibited continuous proliferation over a period of 4 days (**Supplemental Figure 3**).

**Figure 2.**
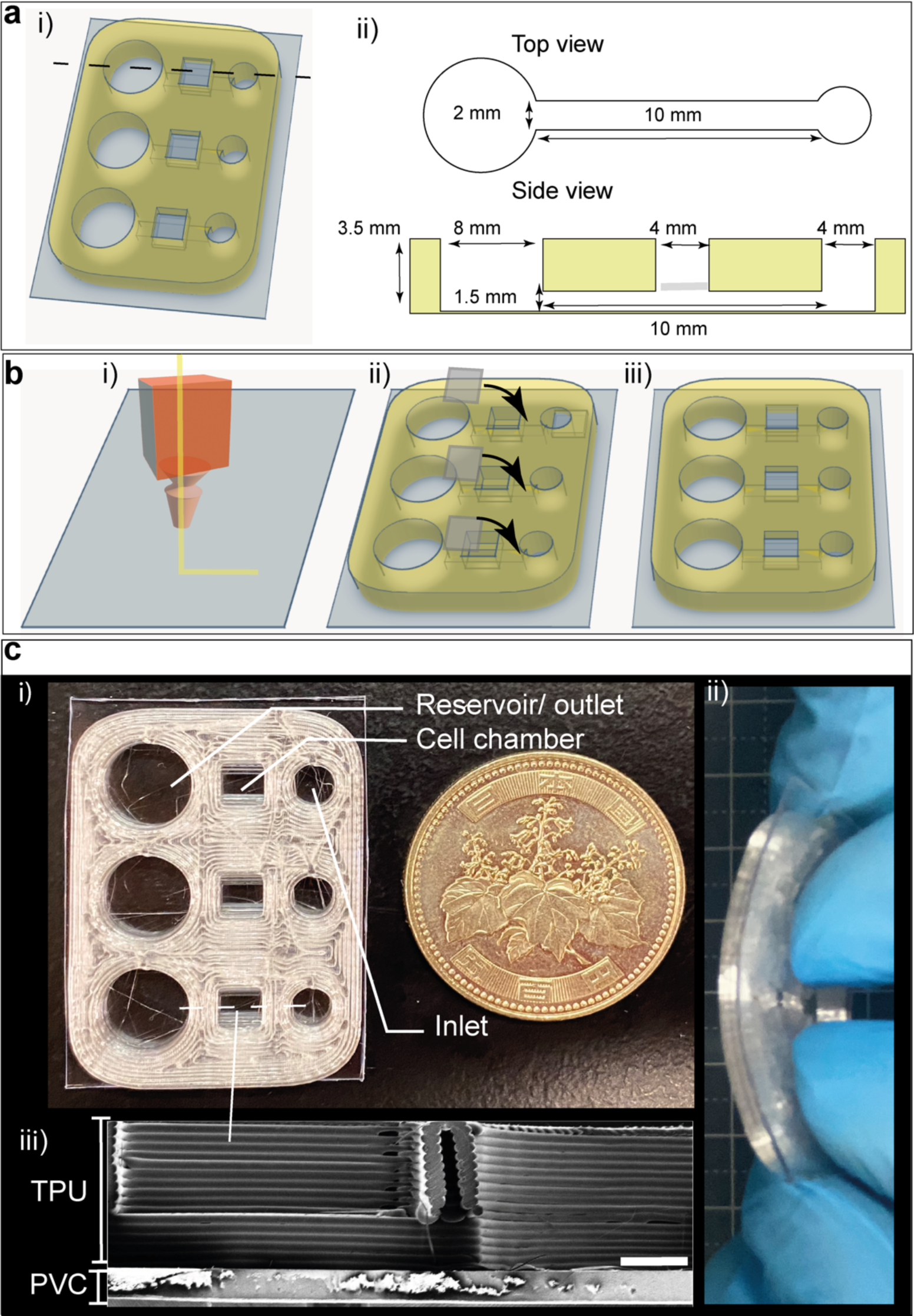
Conceptual design and features of the 3D-FlexTPU-MFD. (a) Detailed design of the cell culture chambers: i): 3D perspective view showing multiple chambers, inlet and outlet ports. ii): Top view and side view of the chamber dimensions. The main chamber has a width of 2 mm and a length of 10 mm, with outlet and inlet segments measuring 8 mm and 4 mm. The side view shows chamber depths of 1.5 mm and overall device thickness of 3.5 mm. (b) Fabrication process of the 3D-FlexTPU-MFD: i) Deposition of TPU polymer onto PVC sheets, followed by sequential 3D printing of microfluidic channels, inlet, and outlet reservoirs. ii) Assembly by overlaying PVC covers onto the printed channels. iii) Final device preparation by trimming the edges. (**c**) i) A photograph of the fully assembled 3D- FlexTPU-MFD. ii) A demonstration of the device’s flexibility through repeated manual twisting. iii) Scanning electron microscopy (SEM) image showing a cross-section of the device, including the cell chamber and inlet bonded to the PVC substrate. Scale bar: 1000 μm.

### 3.2. Myoblast differentiation and maturation in the 3D-FlexTPU-MFD

The differentiation and alignment of myoblasts within the 3D-flexTPU-MFD was assessed on days 3 (D3), 6 (D6), and 9 (D9) using Calcein AM and Hoechst 33342 staining (**Figure 3a**). Over time, cells exhibited increased elongation and alignment, with pronounced formation of organized myotube-like structures observed from D6. Quantitative analysis revealed a significant increase in fluorescence intensity and myotube area from D3 to D9, highlighting cells survival in the device (**Figure 3b**). To evaluate the impact of the 3D-flexTPU-MFD on cellular morphology, F-actin staining was used to compare cells alignment on D3 and D6 with that of cells cultured in a 96-well plate at D6 (**Figure 3c**). Cells in the 3D device exhibited superior alignment and elongation, while cells in the 96-well plate displayed less organized structures. Myotube width, a key indicator of differentiation quality, was measured and compared across different culture conditions (**Figure 3d**). Myotubes cultured in the device on Day 3 (D3) were significantly wider than those on Day 6 (D6) and those cultured in the 96-well plate. However, no significant difference was observed in myotube widths between cells cultured in the device and those in the 96-well plate on D6, confirming the successful formation of myotubes under both conditions. Further, angular analysis of myotube orientation revealed a higher degree of alignment in the 3D-flexTPU-MFD compared to the 96-well plate (**Figure 3e**). The narrower angular distribution of myotubes in the device reflects its ability to mimic the aligned architecture of native skeletal muscle tissue.

**Figure 3.**
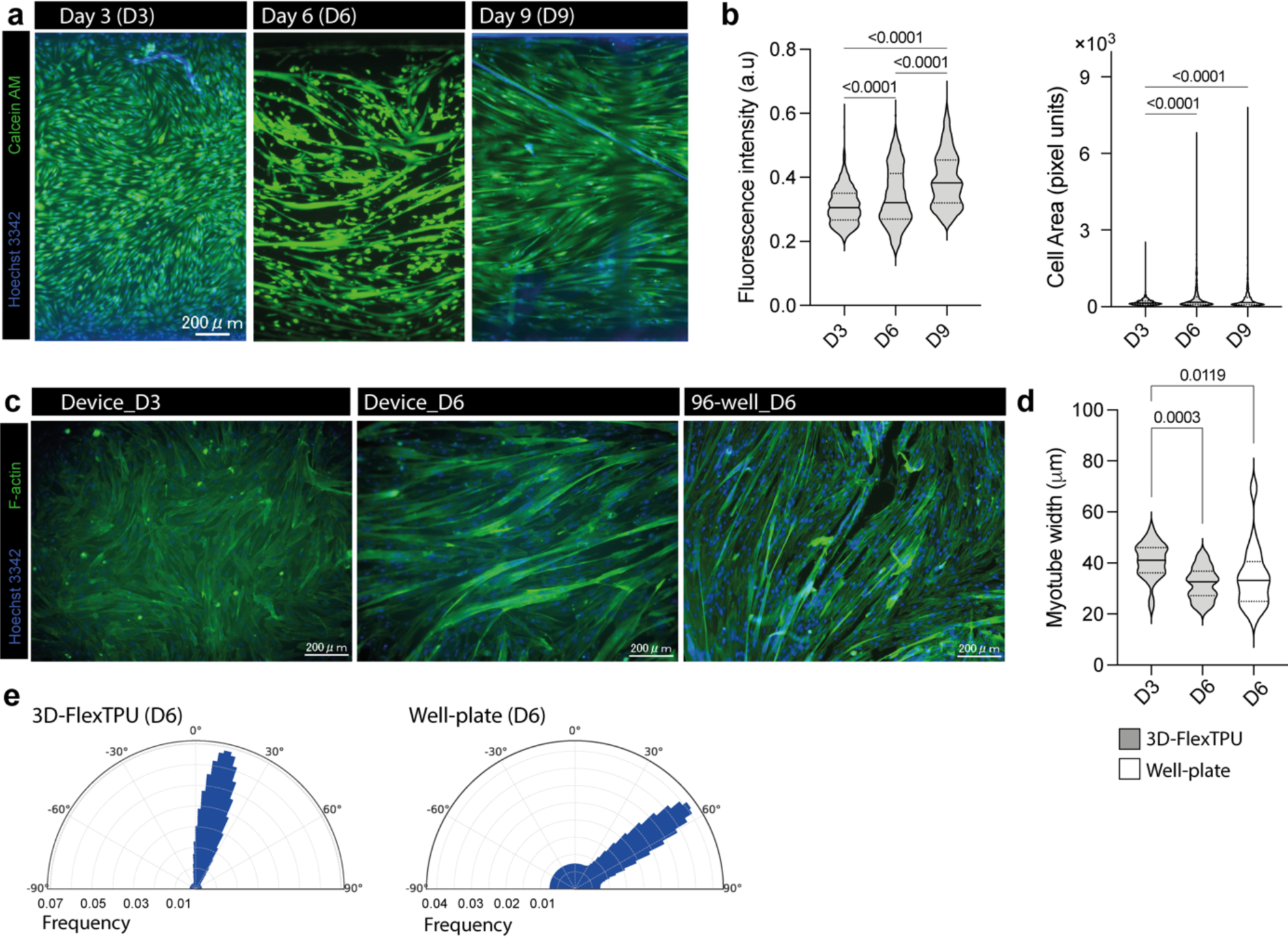
Evaluation of myoblast differentiation, alignment, and maturation in the 3D- flexTPU-MFD. (a) Representative images of myoblasts cultured in the 3D-flexTPU-MFD on day 3 (D3), day 6 (D6), and day 9 (D9), stained with Calcein AM (green) to assess cell viability and Hoechst 33342 (blue) to visualize nuclei. Scale bar: 200 µm. (b) Quantitative analysis of fluorescence intensity (left) and cell area (right) from myotube structures on D3, D6, and D9. (c) F-actin staining of myoblasts cultured in the 3D-flexTPU-MFD on D3 and D6 and in a 96-well plate on D6. Scale bar: 200 µm. (d) Violin plots comparing myotube widths in the 3D-flexTPU-MFD at D3 and D6 and in the 96-well plate at D6. (e) Polar histograms representing the angular distribution of myotubes cultured in the 3D-flexTPU-MFD (left) and the 96-well plate (right) on D6.

### 3.3. Attachment and neurite outgrowth of human iPSC-derived optic vesicle organoids in the 3D-FlexTPU-MFD

To evaluate the compatibility of the 3D-FlexTPU-MFD with human iPSC-derived cells, we generated iPSC-derived optic vesicle (OV) organoids and cultured them within the devices. We further assessed neurite outgrowth, cell viability, and marker expression. **Figure 4a** shows phase-contrast images of organoids on Day 1 (D1) and Day 3 (D3). On D1, minimal neurite outgrowth was observed. By D3, significant neurite extension had occurred. Quantitative analysis revealed a marked increase in neurite length, with a statistically significant difference between D1 and D3. Cell viability within the organoids was assessed on Day 2 (D2) using Calcein AM (green) and Hoechst 3342 (blue) staining (**Figure 4b**). The images showed widespread Calcein AM staining, indicating high cell viability and structural integrity. Notably, viable cells with distinct neuronal morphology were observed at the periphery of the organoids, confirming that the microfluidic device provides a favorable environment for neuronal growth and differentiation (**Figure 4b, ii**). To further evaluate organoid differentiation, we examined the expression of the eye developmental marker PAX6 on Days 3 (D3) and 6 (D6). Fluorescence images for Hoechst 3342 (blue) and PAX6 (yellow) demonstrated consistent expression of PAX6-positive cells on both days (**Figure 4c, i and ii**). 3D reconstructions confirmed the widespread distribution of PAX6-expressing cells within the organoids (**Figure 4c, iii**).

**Figure 4.**
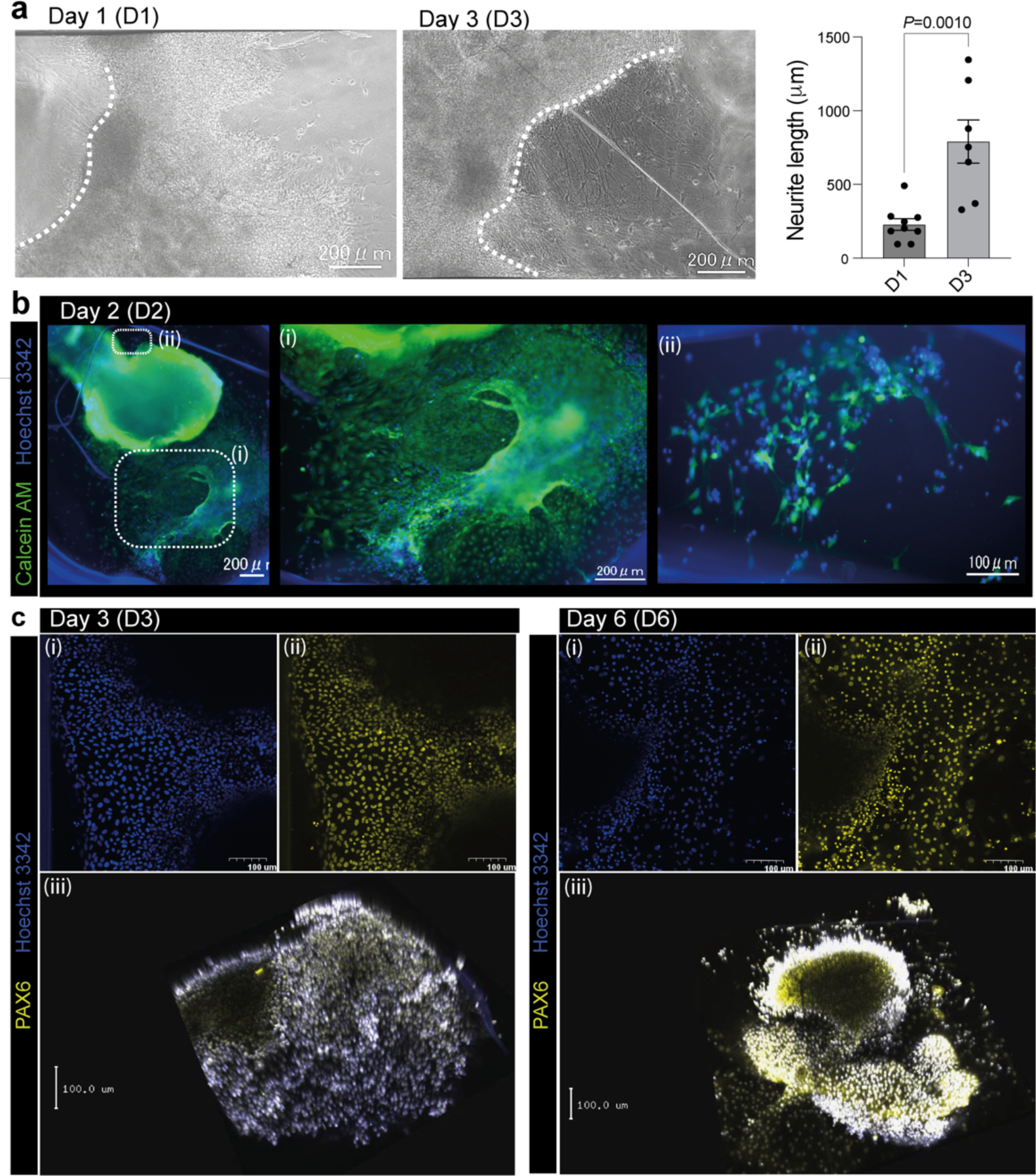
Attachment and differentiation of iPSC-derived optic vesicle organoids in the 3D-FlexTPU-MFD. (a) Brightfield images of iPSC-derived optic vesicle organoids cultured in the 3D-FlexTPU- MFD at day 1 (D1) and day 3 (D3). Dashed lines mark the edges of the organoids, highlighting enhanced attachment and neurite outgrowth over time. Quantification of neurite length shows a significant increase from D1 to D3 (*P = 0.0010*). Scale bar: 200 μm. (b) Fluorescence images of organoids at day 2 (D2) stained with Calcein AM (green) and Hoechst 33342 (blue). (i) and (ii) illustrate viable cell populations and initial attachment regions. Scale bars: 200 μm (i) and 100 μm (ii). (c) Immunofluorescence staining of cells at day 3 (D3) and day 6 (D6), showing the expression of PAX6 (yellow), an eye developmental marker and nucleus staining with Hoechst 33342 (blue) and 3D reconstructions using confocal scanning microscopy (iii). Scale bars: 100 μm.

## 4. Discussion

The development of fast, simple, and cost-effective methods for constructing OoC models is essential for advancing alternative approaches in drug discovery and disease modeling. In this study, we demonstrate that the use of thermoplastic TPU and PVC allows for the one-step fabrication of flexible microfluidic devices through FDM-based 3D printing.

We selected TPU due to its favorable characteristics, such as elasticity, optical clarity, and suitable hydrophobicity^14^. The water contact angle of TPU is 110° ^15^, which is comparable to that of polydimethylsiloxane (PDMS) that has a water contact angel of 107° ^16^. This property allows TPU to absorb moisture and facilitate gas exchange, enhancing cell viability. Additionally, PVC has a water contact angle of 99.5° ^17^, making it hydrophobic, which can support the bonding process with TPU through possible hydrophobic interactions. In contrast, other thermoplastics such as polylactic acid (PLA) and acrylonitrile butadiene styrene (ABS) showed very low bonding affinity to PVC substrates as compared with TPU (data not shown) possibly due to their hydrophilic nature^14^. TPU, however, formed a strong bond with PVC that remained intact under mechanical stresses, such as pulling or bending, demonstrating the feasibility and robustness of the fabrication process.

To determine the optimal conditions for fabricating a microfluidic device using TPU and PVC suitable for culturing mammalian cells, we designed various channel geometries, including a letter “R,” an array of channels with different widths, and a serpentine channel. While all these structures were printed with good fidelity, the channel array revealed that channels with widths of 0.5 mm or less exhibited lower printing fidelity ratios. This reduced fidelity can likely be attributed to the use of a 0.4 mm extrusion head, which makes printing narrower channels more challenging.

Based on these findings, we designed a microfluidic device_3D-flexTPU-MFD_ featuring three cell culture chambers, fabricated in a single step using 3D printing. Each chamber had a channel width of 2 mm, a length of 10 mm, and a height of 1.5 mm. Additionally, the cell culture channels were connected to an inlet on one side and an outlet reservoir on the other, ensuring a continuous supply of nutrients and facilitating the collection of metabolites. We tested 3D-flexTPU-MFD by culturing human primary myoblasts and evaluating their differentiation into myotubes over a 9-day period., alignment, and structural organization compared to conventional 2D culture systems. Interestingly, myotubes cultured in the 3D- flexTPU-MFD exhibited a parallel orientation, while those grown in well plates showed a more random orientation. This suggests that the fabricated device not only supports myoblast growth and differentiation but also enhances the morphological alignment of cells compared to traditional culture systems.

Human iPSC-derived organoids are powerful tools for disease modeling and drug testing^18^. In this study, we generated iPSC-derived optic vesicle organoids a transient stage of cells before their differentiation into forebrain or retinal organoids using a modified version of our previously established method^19,20^. On day 20, the organoids were cultured within the 3D-flex TPU-MFD, and cell viability was confirmed through Calcein AM staining. Notably, the organoids exhibited progressive neurite outgrowth, indicating the functional differentiation of neuronal cells within the organoids. Furthermore, the organoids maintained high levels of the eye and forebrain marker PAX6 at days 3 and 6, supporting the preservation of cell identity and functionality. These results indicate that the 3D-FlexTPU-MFD supports the maintenance of human iPSC derived cells differentiation during culture and promotes neurite outgrowth, maintains high cell viability, and supports the expression of key eye developemtnal markers, making it a promising platform for organoid-based models in disease modeling research and drug testing.

While this study successfully demonstrated the feasibility of using the 3D-FlexTPU-MFD for cell culture of cells in 2D and 3D, one limitation is that experiments were conducted under static conditions. Incorporating flow perfusion systems in future work could improve nutrient distribution and more closely mimic physiological environments. Nevertheless, the current static setup effectively supported cell viability, differentiation, and alignment, highlighting the device’s robustness. Future studies could expand on these promising results by exploring applications in drug testing and disease modeling.

## 5. Conclusion

In this study, we developed 3D-FlexTPU-MFD using a one-step FDM process with TPU as the filament and PVC as the bonding substrate. The device supported the culture of human primary myoblasts, endothelial cells, and iPSC-derived OV organoids. Myoblasts exhibited high viability and formed aligned myotube bundles, outperforming conventional well-plate cultures. iPSC-derived OV organoids-maintained viability, formed neurites, and expressed the forebrain and eye marker PAX6. The 3D-FlexTPU-MFD offers a versatile platform for cell culture, tissue modeling, and organ-on-a-chip applications. Future work will explore flow perfusion systems and applications in drug testing and disease modeling.

## Authorship contribution

Rodi Kado Abdalkader was responsible for the overall research concept, project management, experimental design and execution of tests, data analysis and interpretation, results visualization, and manuscript writing. Satoshi Konishi and Takuya Fujita provided resources and contributed to manuscript refinement. All authors have reviewed and approved the final version of the manuscript for publication.

## Acknowledgments

We acknowledge the Ritsumeikan Global Innovation Research Organization (R-GIRO) for their support. We are also gratefully acknowledging the Biomedical Engineering Center (BMEC) at Ritsumeikan University for providing access to the confocal scanning microscopy facility.

## Supplementary Information

**Supplementary Figure 1.**
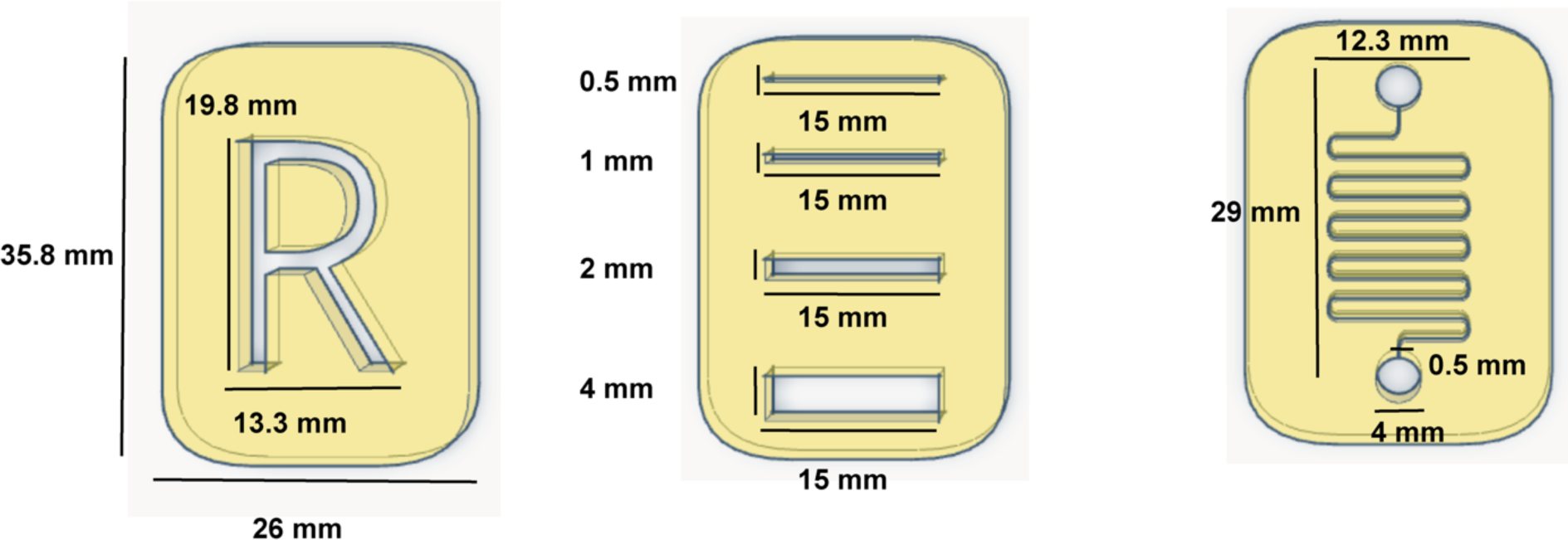
Design specifications of microfluidic devices. Schematic diagrams of various channel designs with dimensions: Left: Letter “R” design (19.8 mm height × 13.3 mm width) within a 35.8 mm × 26 mm rectangular base. Middle: Three rectangular straight channels, each 15 mm in length, with widths of 0.5 mm, 1 mm, 2 mm, and 4 mm. Right: Serpentine channel design measuring 29 mm × 12.3 mm, with a channel width of 0.5 mm and inlet width of 4 mm.

**Supplementary Figure 2.**
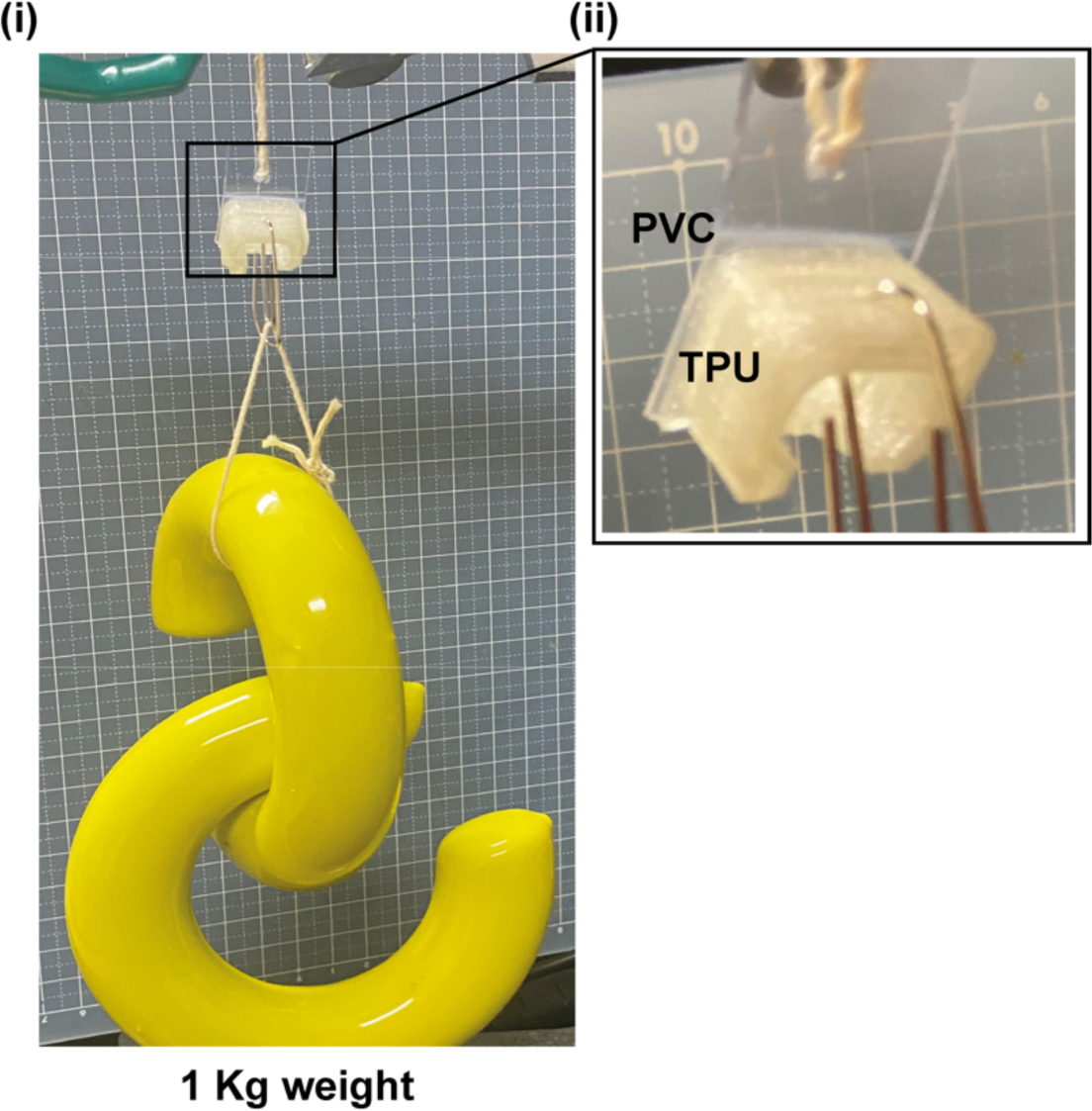
Mechanical stability of the TPU-PVC construct under load. (i) The TPU-PVC construct suspending a 1 kg weight, demonstrating its mechanical stability and robustness without visible deformation or failure. (ii) A close-up image of the TPU-PVC interface, showing the strong integration of the two materials.

**Supplementary Figure 3.**
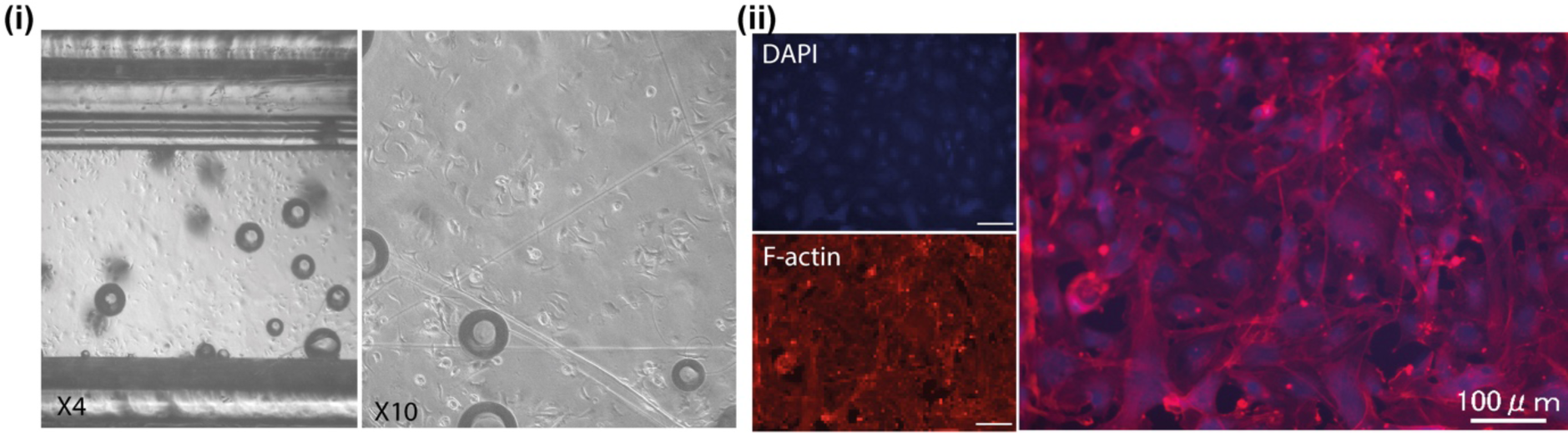
Culturing and adherence of HUVECs in the 3D-FlexTPU-MFD. (i) Brightfield microscopy images of HUVECs at 3 hours of culture within the microfluidic device, showing initial cell attachment and distribution. (ii) Staining of HUVECs at day 4 of culture, highlighting the cytoskeletal organization with F-actin (red) and nuclear localization with DAPI (blue). Scale bars of 100 μm.

